# Targetable Tetrazine-Based Dynamic Nuclear Polarization Agents for Biological Systems

**DOI:** 10.1101/740530

**Authors:** Byung Joon Lim, Bryce E. Ackermann, Galia T. Debelouchina

**Affiliations:** Department of Chemistry and Biochemistry, University of California, San Diego, La Jolla, CA 92093, USA

## Abstract

Dynamic nuclear polarization (DNP) has shown great promise as a tool to enhance the nuclear magnetic resonance (NMR) signals of proteins in the cellular environment. As the sensitivity increases, the ability to select and efficiently polarize a specific macromolecule over the cellular background has become desirable. Here, we address this need and present a tetrazine-based DNP polarization agent that can be targeted selectively to proteins containing the unnatural amino acid (UAA) norbornene-lysine. The UAA can be introduced efficiently by genetic means in the cellular milieu. Our approach is bio-orthogonal and easily adaptable to any protein of interest. We illustrate the scope of our methodology and investigate the DNP polarization transfer mechanisms in several biological systems. Our results present the first molecular view of the complex polarization transfer pathways in targeted DNP and ultimately pave the way to selective DNP-enhanced NMR spectroscopy in both bacterial and mammalian cells.

---

Dynamic nuclear polarization (DNP) has made significant impact on the solid-state nuclear magnetic resonance (NMR) investigations of complex biological systems.^1–2^ With now routine signal enhancements in the 100-fold range, DNP has aided the *in vitro* structural studies of amyloid polymers,^3–4^ membrane proteins^5–6^ and biological material such as collagen, bones and tissues^7–8^. Improvements in DNP instrumentation, sample preparation and polarization agent design have also allowed NMR spectroscopists to turn their attention to the cellular environment where the structural analysis of endogenous concentrations of biological macromolecules has come within reach.^3, 9–11^ In a typical DNP experiment, the sample is doped with millimolar amounts of polarization agents, small molecules that contain two stable unpaired electron spins. The sample is cryoprotected, frozen to ∼ 100 K and subjected to continuous microwave irradiation during magic angle spinning (MAS). The generated electron spin polarization is transferred to the nuclear spins, resulting in significant enhancement of the NMR signals. Concurrently, multidimensional NMR experiments are performed and structural constraints obtained with much improved sensitivity. While this strategy has been applied to most *in vitro* and cellular DNP experiments to date, it suffers from one major limitation: the polarization transfer is non-selective and results in enhancements for all molecules in the sample, including the cellular background.^10–11^ To overcome this shortcoming, several biochemical strategies can be employed including overexpression of the protein of interest^12–13^ or the introduction of recombinant isotopically labeled protein into cells by electroporation^11^. These approaches, however, require extensive perturbation of the cellular milieu and are not applicable to the structural studies of endogenous concentrations of cellular components. Alternatively, the design of targeted DNP polarization agents has been explored with selectivity achieved either through cysteine chemistry or protein-ligand interactions.^14–20^ The applications of these strategies, however, are limited due to the reducing cellular environment or the requirement for a known synthetically accessible ligand of the target protein. We, therefore, sought to develop a general targeting strategy that can be applied to any protein of interest in a cysteine-independent manner, and that is compatible with DNP-enhanced NMR studies of proteins in the cellular interior.

Our targeting approach is based on the bio-orthogonal chemical reactions between tetrazines and strained cycloalkenes.^21–22^ These reactions display superb efficiency and selectivity in the cellular milieu and are routinely used to label cellular proteins with optical probes.^23–26^ Here, we harness their beneficial properties to attach DNP polarization agents to a variety of proteins (Fig. 1a). To this end, we started by coupling a tetrazine moiety to TOTAPOL, a stable, nitroxide-based DNP polarization agent^27^ (**SI Fig.1**). The TOTAPOL-tetrazine biradical (TTz) displays the characteristic nitroxide hyperfine splitting in its EPR spectra and can be efficiently conjugated to strained cycloalkenes such as norbornene. Next, we synthesized the unnatural amino acid (UAA) norbornene-lysine and introduced it at position 6 in the sequence of the model protein ubiquitin through amber suppression (Fig. 1b**; SI Fig. 2)**. Amber suppression is a molecular biology technique that enables the introduction of UAAs in living cells by genetic means through the reassignment of the amber (TAG) stop codon.^28^ The UAA is incorporated at the desired TAG position through an orthogonal tRNA and aminoacyl-tRNA synthetase (aaRS) pair introduced into the cell via a separate plasmid construct. Our application required that we perform amber suppression in M9 media to ensure the ^13^C,^15^N-labeling of the targeted protein.

**Figure 1.**
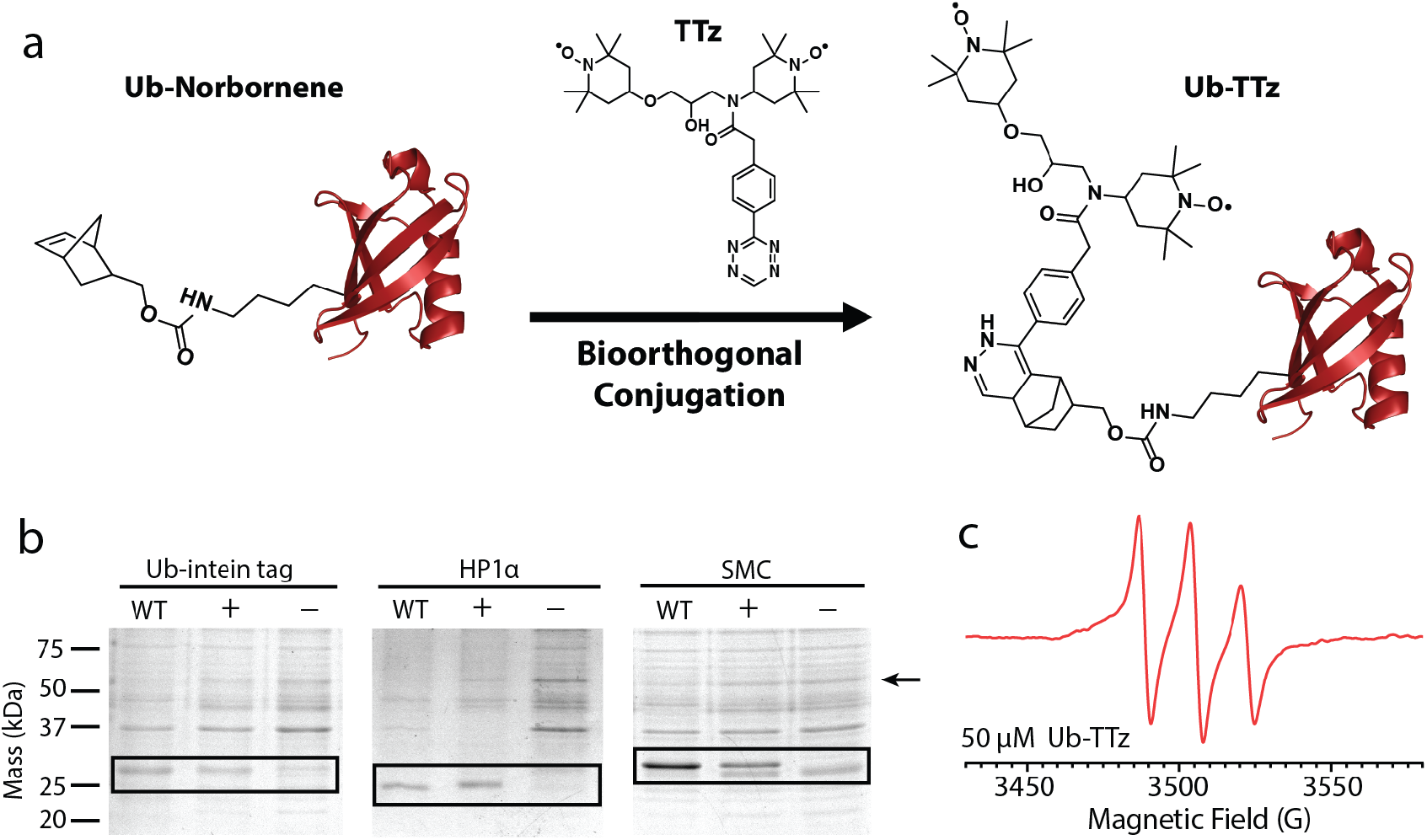
a) A general polarization agent targeting strategy based on the bio-orthogonal reaction between tetrazine and norbornene. b) Incorporation of norbornene-lysine into three different proteins by amber suppression in *E. coli* grown in M9 media supplemented with ^13^C and ^15^N-labeled components. c) 9 GHz EPR spectrum of 50 μM ubiquitin-TTz.

After testing several plasmid vectors and constructs, we chose the pULTRA vector where the expression of the tRNA/aaRS pair is under the control of a *lacI* promoter^29^ and we combined it with a tRNA/aaRS sequence optimized for the incorporation of norbornene-lysine^24^. Using this strategy, we observed high yields of isotopically labeled and modified proteins, not only for ubiquitin but also for several other systems of varying size and complexity including heterochromatin protein 1 (HP1α) and the structural maintenance of chromosomes (SMC) protein (Fig. 1b). Armed with these reagents, we next performed the biorthogonal reaction to produce protein-TTz conjugates. The reaction between tetrazine and norbornene was robust, with high efficiency under both denaturing and native conditions. After successful protein-TTz conjugation, excess TTz was removed by dialysis and the expected 1:1 ratio of protein to TTz was confirmed by LC-MS and EPR (Fig. 1c**, SI Fig. 3 & 4**).

We next proceeded to determine the efficiency of TTz as a polarization source in comparison to the standard DNP sample preparation where the excess polarization agent is evenly distributed throughout a glassy matrix. As a direct measure of the enhancement, we used the signal in the presence of microwaves divided by the signal in the absence of irradiation. Under these conditions, we measured an enhancement of 24 for 1 mM ubiquitin-TTz which is 68% of the enhancement provided by dispersed TOTAPOL despite the 15-fold difference in radical concentration (Fig. 2a). The enhancement observed in our conjugated system is comparable or higher than the enhancements reported for other targeted-DNP approaches^14^ and it was sufficient to record a 2D ^13^C-^13^C spectrum of just 100 ug of ubiquitin (1 mM) in less than a day (Fig. 2b). This spectrum presents many resolved correlations similar to the recently published and partially assigned DNP spectra of ubiquitin samples prepared with dispersed AMUPol.^11^

**Figure 2.**
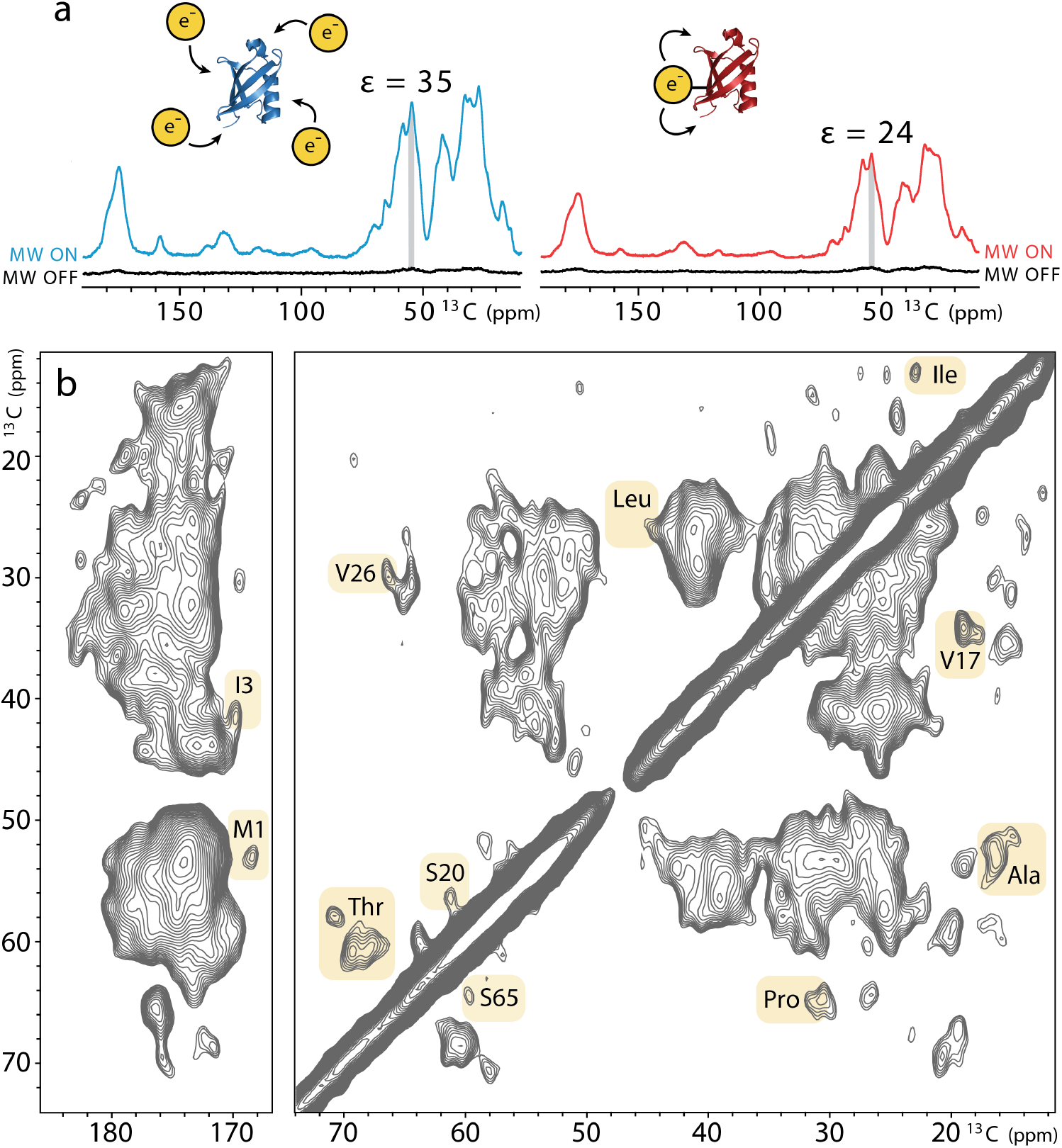
a) Comparison of the DNP enhancements measured in a sample containing 1 mM wild-type ubiquitin with 15 mM dispersed TOTAPOL (left) and 1 mM ubiquitin-TTz (right). Spectra were acquired at 600 MHz ^1^H Larmor frequency, 12 kHz MAS, 100 K. b) DNP-enhanced 2D ^13^C-^13^C CORD correlation^30^ spectrum of 1 mM ubiquitin-TTz acquired at 24 kHz.

DNP enhancements depend on many experimental parameters including polarization agent concentration, interscan delay and MAS frequency.^1, 27, 31–32^ These parameters can provide valuable information regarding the DNP polarization transfer pathways, yet, have not been extensively investigated in targeted systems similar to the ones presented here. Therefore, we compared conjugated samples at 1 mM protein (i.e. 1 mM TTz) and dilute samples at ∼ 50 μM protein and TTz (Fig. 3). The measured enhancements at lower concentration were smaller, in the 6 – 10 range for different proteins. It should be noted, however, that dispersed TOTAPOL or TTz samples with similar concentration of biradicals yielded no enhancement. Therefore, conjugation significantly improves DNP polarization transfers by bringing the polarization source and target in close proximity to each other. The amount of protein in these samples is in the 5 - 10 μg range and is close to the endogenous amounts of protein present in 10^6^ mammalian cells that can easily fit into an NMR rotor.^11, 33^ The targeted proteins also vary in shape, size and sequence, where ubiquitin is small and globular^34^, HP1α is a flexible homodimer^35^, and SMC is a coiled-coil protein with an extreme length of 17 nm^36^. The small differences in enhancement among these systems may be due to the characteristics of the protein sequence, the overall structure of the protein or the choice of conjugation site.

**Figure 3.**
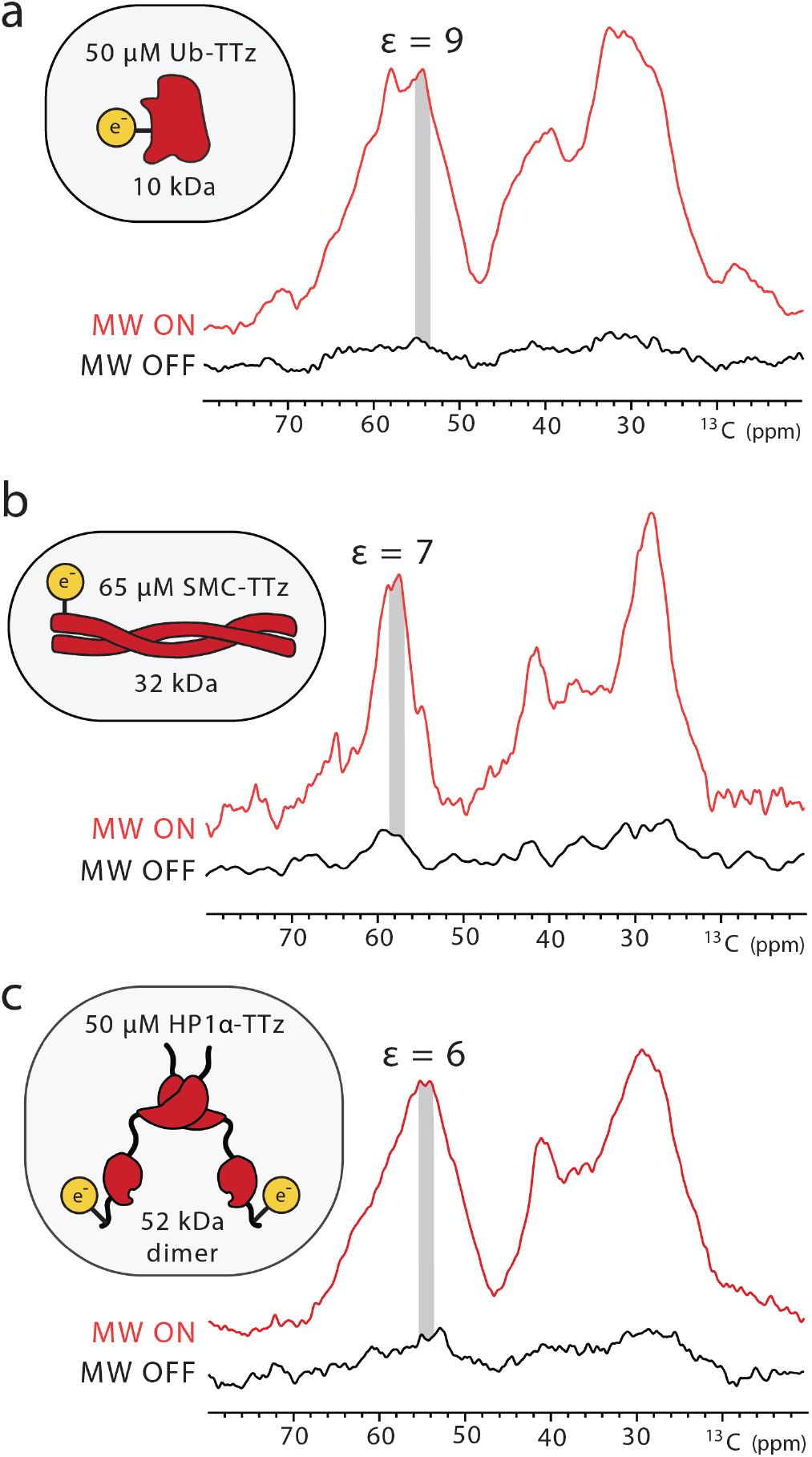
DNP enhancements for low concentrations of a) Ub-TTz; b) SMC-TTz; and c) HP1α-TTz.

Intrigued by the concentration dependence of the enhancements in our targeted systems, we also measured polarization buildup curves that report on the distance dynamics of polarization transfers (Fig. 4**, SI Fig.5**).^37–38^ To this end, we compared the buildup behavior of a sample prepared with 15 mM dispersed TOTAPOL and samples containing 1 mM or 50 μM Ub-TTz. While the dispersed sample displayed a monoexponential buildup curve with a 5.8 s time constant, the behavior of the Ub-TTz samples appeared more complex. In particular, the buildup curve for the 50 μM Ub-TTz sample is consistent with two events, one with a short buildup time, and another event characterized with a much longer time constant. Considering the low concentration of Ub-TTz in this sample, we interpret the first buildup event as the fast polarization transfer from TTz to the directly conjugated protein (∼ 3 nm in size), while the second step reflects long distance transfer from one conjugate to another (∼ 50 nm distance). Furthermore, our observations also suggest that at ∼ 1 mM Ub-TTz where the distance between biradicals is ∼ 15 nm, the buildup curve is at the turning point between mono- and bi-exponential behavior. In this case, each protein is polarized by its own source as well as TTz moieties on neighboring proteins, resulting in larger enhancements and buildup times on the order of 8-12 s. It should be noted that almost all published targeted DNP studies so far have been performed at concentrations of 1 mM or above, and, thus, most likely, the reported enhancements reflect a significant intermolecular polarization transfer component.

**Figure 4.**
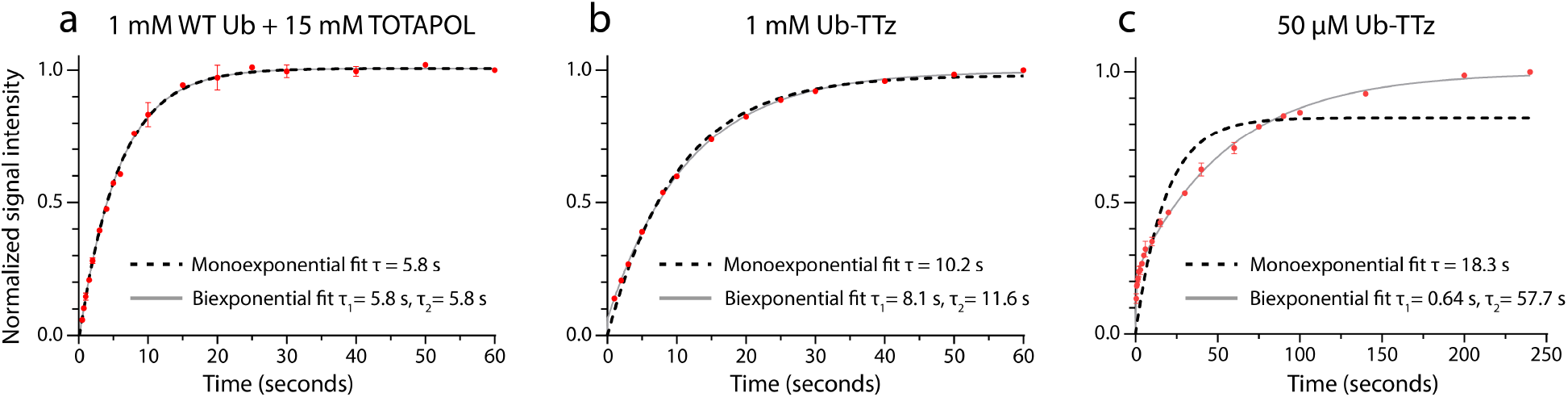
Polarization buildup curves for a) 1 mM wild-type ubiquitin with 15 mM dispersed TOTAPOL; b) 1 mM Ub-TTz; and c) 50 μM Ub-TTz.

**Figure 5.**
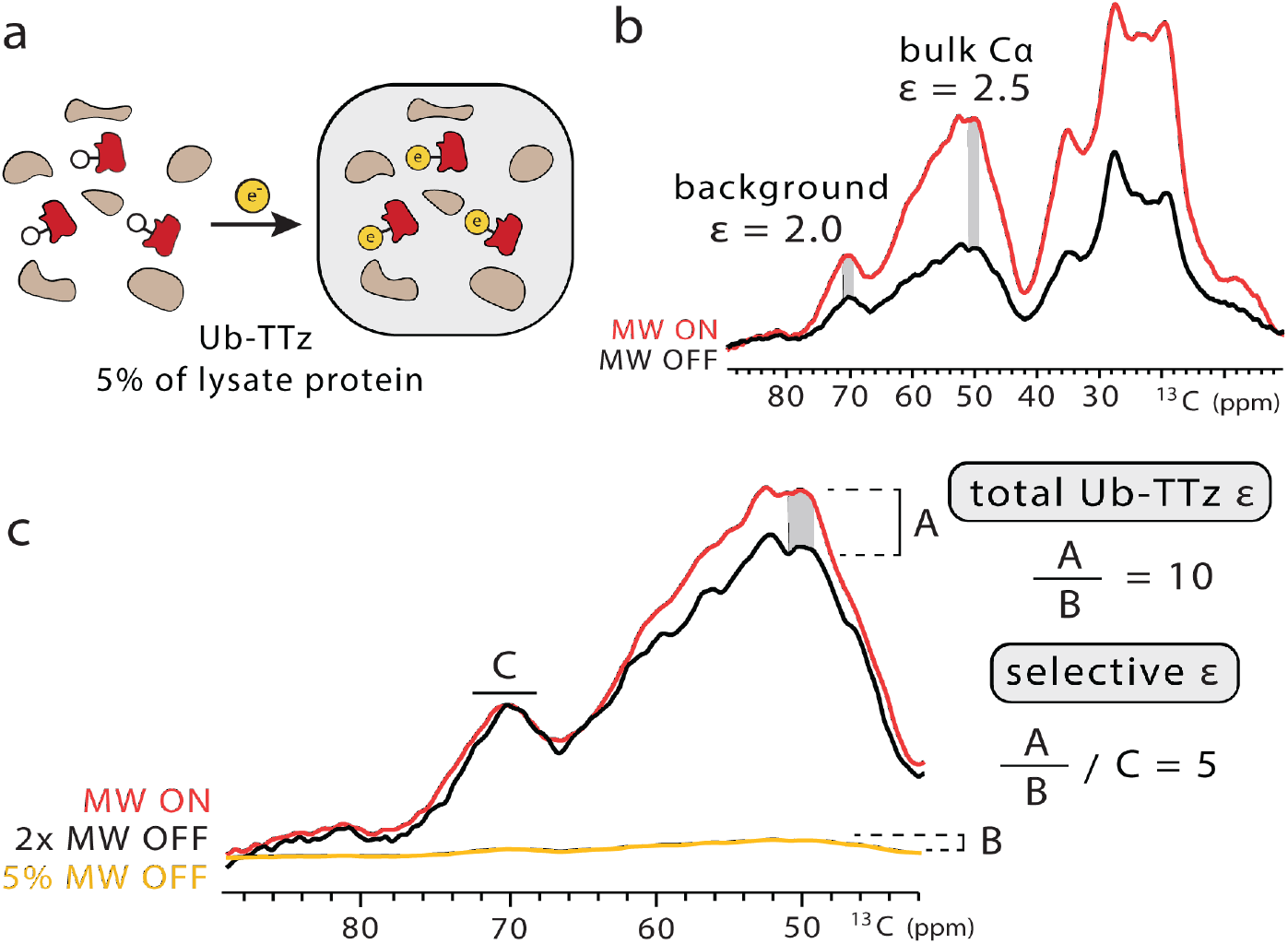
a) Targeted DNP strategy for bacterial lysates. b) Overall signal enhancement of the ^13^C-labeled lysate. c) Strategy to determine the selective enhancement of Ub-TTz in the lysate. The microwave (MW) off signal was scaled to the microwave on signal so that the glycerol solvent peaks matched in intensity. The leftover enhancement was denoted as *A*. Then, the microwave off signal was scaled down to reflect the 5% abundance of Ub-TTz in the sample and the intensity of this rescaled spectrum was denoted as *B*. The ratio of *A* to *B* produced the total enhancement on ubiquitin. The selective enhancement is equal to the total ubiquitin enhancement divided by the background enhancement.

Finally, we explored the effects of MAS frequency (**SI Fig. 6 & 7**). Previous studies have reported an unfavorable relationship between MAS frequency and DNP enhancements due to a complex time dependent modulation of the electron and nuclear dipolar couplings.^31, 39–41^ As expected, increasing the MAS frequency from 12 kHz to 24 kHz reduced the enhancements in both dispersed and conjugated samples although the reduction was much less pronounced for Ub-TTz at low concentration. For example, the signal intensities for the Cα region decreased by 44% for the dispersed sample, 49% for the 1 mM Ub-TTz sample and 15% for the 50 μM Ub-TTz sample. These results suggest that tethering the biradical to the protein of interest suppresses the unfavorable MAS relationship, presumably due to a less pronounced dependence on long-distance polarization transfers that may be affected by MAS. Therefore, targeted DNP might be better suited for experiments under fast MAS, an imminent and exciting goal for the DNP NMR community.

To further demonstrate the scope of our targeted bio-orthogonal approach, we extended our studies to a more complex setting. In particular, we targeted isotopically labeled bacterial lysates containing overexpressed ubiquitin with the norbornene-lysine UAA. After incubation with TTz, we dialyzed the lysate to remove unreacted TTz and confirmed complete TTz to protein conjugation by LC-MS (**SI Fig. 8**). We performed biochemical assays to estimate the ubiquitin amount in this sample, which was ∼ 5% of the total protein and was present at ∼ 40 μM in the DNP rotor (**SI Fig. 8**). Using the background subtraction approach introduced by Viennet et al.^18^, we estimate that the selective enhancement for conjugated ubiquitin is 5-fold greater than the enhancement of the lysate background. Our overall ubiquitin enhancement in this setting is ∼10, which is on par with the enhancements we measured for similar concentrations of purified protein-TTz conjugates. The ratio of specific to background enhancement in lysates can be tuned by the interscan delay, where shorter delays favor the targeted protein due to the close proximity of the paramagnetic species (**SI Fig. 9**). The specificity can be improved even further by coupling targeted DNP with selective isotopic labeling strategies and direct protein expression where only the protein of interest is isotopically labeled and enhanced.^42^

In summary, our tetrazine-based agents represent a general and efficient strategy to target DNP polarization to a variety of protein systems both *in vitro* and in complex biological settings. Since tools for amber suppression exist for different organisms, our approach is applicable to virtually any protein in bacterial or mammalian cells where tetrazine-based reagents have exhibited good permeability and superb reactivity and selectivity.^23, 26, 43^ In this study, we chose to couple tetrazine to TOTAPOL due to the ease of synthesis of the biradical, however, our strategy is compatible with other improved DNP polarization agents such as AMUPol.^44^ Our results also demonstrate that much is still to be learned about the polarization transfer mechanisms in targeted DNP. For the first time, we show that at low protein concentrations, the intramolecular short-range polarization transfer can be decoupled from the intermolecular long-range transfer mechanisms. We also explore the relationship between enhancement and MAS frequency and expect that targeted DNP will outperform conventional DNP strategies as improvements in instrumentation bring DNP into the fast MAS regime. A thorough understanding of the DNP polarization transfer mechanisms in conjugated systems will be essential in the selective signal enhancement of proteins of various sizes and shapes and on preventing polarization “leakage” into the cellular background. We are also excited at the prospect of combining our targeted DNP approach with other developments in the field such as higher magnetic fields,^45–46^ time domain DNP experiments^47^ and optical approaches^10^ for the comprehensive description of protein structures in cells.

## Acknowledgements

We would like to thank Dr. Ivan Sergeyev at Bruker Biospin and Justin Lee in the Borovik lab at UCI for help with EPR data acquisition, Dr. Peter Schultz and Dr. Minseob Koh for advice regarding amber suppression, Dr. Carsten Schultz and Dr. Jan-Erik Hoffmann for the aaRS construct, Brysa Alvarado and Melinda Serrato for assistance with TOTAPOL synthesis, and members of the Debelouchina and Opella labs for helpful discussions. B. A. was supported by NIH Molecular Biophysics Training Grant T32 GM008326 and B. J. L. and G. T. D. were supported by UCSD startup funds. This work utilized the Biotechnology Research Center for NMR Molecular Imaging of Proteins at UCSD, supported by NIH grant P41 EB002031.

