## Supplementary methods and figures for "Targetable Tetrazine-Based Dynamic Nuclear Polarization Agents for Biological Systems"

#### Materials and equipment

All reagents and solvents were purchased from Sigma Aldrich (St. Louis, MO), Thermo-Fisher Scientific (including Fisher Scientific and Acros Organics, Waltham, MA) and used without further purification unless otherwise noted. Isotopically enriched reagents and solvents for NMR were purchased from Cambridge Isotope Laboratories (Tewksbury, MA). Thin layer chromatography analyses were carried out using aluminum-backed, precoated silica gel plates (Merck TLC silica gel 60 F254) from EMD Millipore (Billerica, MA). Column chromatography was performed using Acros Organics Silica gel (0.035-0.070 mm, 60 Å). $^{1}H$ and $^{13}C$ NMR of small molecules were performed on a Varian Vx 500 MHz spectrometer. Primer synthesis and DNA sequencing were performed by Integrated DNA Technologies (Coralville, IA) and Genewiz (South Plainfield, NJ), respectively. PCR was performed using a Bio-Rad T100 thermocycler (Hercules, CA), while purification of the DNA relied on kits from Biomiga (San Diego, CA) and New England BioLabs (Ipswich, MA). Dialysis kits were purchased from Thermo Fisher Scientific and protein concentrators were obtained from Sartorius (Goettingen, Germany). Reverse-phase (RP) HPLC was performed on a 2545 Binary Gradient Module Waters system equipped with a 2484 UV/vis detector from Waters Corporation (Milford, MA). For prep-scale RP-HPLC purification, we relied on a Waters XBridge BEH C18 19 mm x 250 mm, 10 µm particle size column, while for analytical measurements we used a Waters Symmetry 300 C18 4.6 mm x 150 mm, 5 µm particle size column. HPLC solvent A contained 100% $H_2O$ + 0.1% trifluoroacetic acid (TFA), while solvent B contained 100% acetonitrile + 0.1 % TFA. HR-MS and LC-ESI-TOF MS analysis was conducted on an Agilent 6230 TOF-MS with Jet Stream ESI. Gel images were acquired using a camera and light box from Fotodyne Incorporated (Hartland, WI). Protein and fluorophore absorbances were measured using a Nanodrop One Spectrophotometer by Thermo-Fisher Scientific.

#### Synthesis of polarization agents and UAAs

*Synthesis of norbornene lysine*

Norbornene lysine was synthesized according to the protocol reported by Plass et al.[1] The final product was analyzed by solution NMR spectroscopy and high-resolution mass spectrometry (HR-

MS). $^1$H NMR (500 MHz, DMSO-$d_6$): 7.15 (0.5H, t, $J$ = 5.5 Hz), 7.10 (0.5H, t, $J$ = 5.5 Hz), 6.16 (0.5H, dd, $J$ = 5.6, 3.0 Hz), 6.11-6.07 (1H, m), 5.92 (0.5H, dd, $J$ = 5.6, 2.8 Hz), 4.02–3.97 (m, 0.5H), 3.87–3.81 (m, 0.5H), 3.69–3.64 (m, 0.5H), 3.51–3.48 (2H, m), 3.00-2.90 (2H, m), 2.82–2.75 (2H, m), 2.65 (0.5H s), 2.33–2.25 (0.5H, m), 1.80–1.66 (3H, m), 1.62–1.53 (0.5H, m), 1.35-1.20 (3.5H, m), 1.10–1.1 (1H, m), 0.45 (0.5H, ddd, $J$ = 11.5, 4.1, 2.6 Hz). $^{13}$C NMR (125 MHz, DMSO-$d_6$): δ 171.24, 156.45, 156.40, 137.40, 136.89, 136.26, 132.28, 67.68, 67.11, 51.96, 49.03, 44.70, 43.49, 43.24, 41.78, 41.15, 38.22, 37.94, 29.77, 29.12, 28.97, 28.76, 21.70. HR-MS (ESI): calculated m/z for $[C_{15}H_{25}N_2O_4]^+$ ($[M+H]^+$) is 297.1736, experimentally determined value is 297.1809.

*Synthesis of TOTAPOL-tetrazine (TTz)*

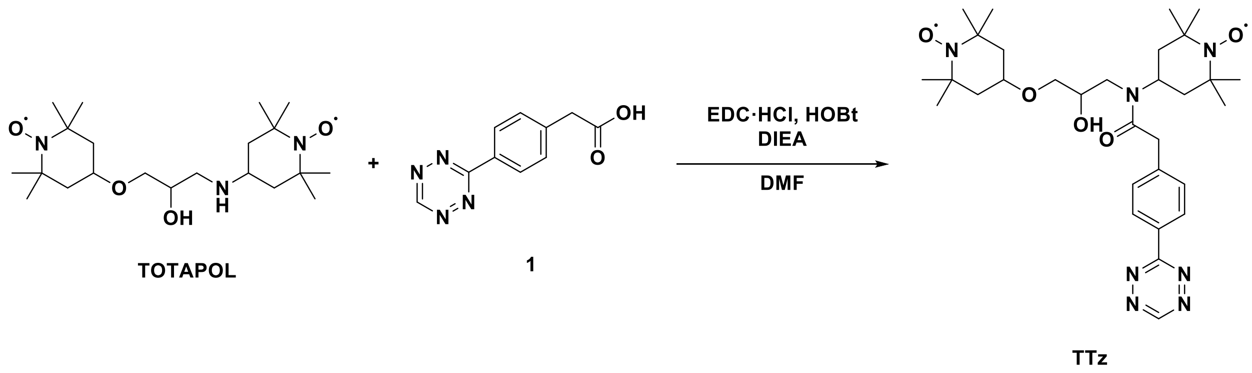


TOTAPOL was synthesized according to the protocol reported by Song et al.[2] Compound **1** was synthesized according to the protocol reported by Yang et al.[3] Compound **1** (100 mg, 0.463 mmol), *N*-(3-dimethylaminopropyl)-*N*'-ethylcarbodiimide hydrochloride (EDC-HCl) (177 mg, 0.925 mmol), 1-hydroxybenzotriazole (HOBt) (125 mg, 0.925 mmol), and *N,N*-diisopropylethylamine (DIEA) (161 mL, 0.925 mmol) were dissolved in dry *N,N*-dimethylformamide (DMF) (5 mL, 0.1 M). TOTAPOL (222 mg, 0.555 mmol) was then added to the solution and the reaction mixture was stirred for 6 hours at room temperature. After completion, the reaction was analyzed by thin layer chromatography (dichloromethane:methanol = 10:1, $R_f$ = 0.38), and ethyl acetate (20 mL) was added to the solution. The organic layer was washed with water (3 times) and brine, then dried over anhydrous sodium sulfate. The crude mixture was filtered, and the solvent was evaporated under reduced pressure. The final product was purified by silica flash column chromatography using ethyl acetate:hexanes (1:2 to 2:1 gradient) as pink solid (yield: 62 %). HR-MS (ESI): calculated m/z for [$C_{31}H_{47}N_7O_5Na^+$] ($[M+Na]^+$) is 620.3531, measured m/z is 620.3525. HPLC trace and EPR spectrum are shown in **SI Fig 1**.

### Preparation of plasmid constructs

#### *pULTRA tRNA$^{Pyl}$/PylRS$^{AF}$ plasmid*

This plasmid encodes the tRNA (tRNA$^{Pyl}$) and the tRNA synthetase (PylRS$^{AF}$) for norbornene lysine incorporation by amber suppression. After initial tests and issues with $^{13}$C label dilution in M9 media, we designed a construct where the pULTRA vector backbone from Chatterjee et al.[4] was combined with the tRNA$^{Pyl}$/PylRS$^{AF}$ insert sequence from Plass et al.[1] The new plasmid was constructed using the NEBuilder® HiFi DNA assembly. Protein expression in this construct is controlled by an IPTG-inducible *lacI* promoter.

Forward (vector): ATGGACAAAAAACCGCTGAA

Reverse (vector): TTACAGGTTCGTAGAGATCC

Forward (PylRS): GGATCTCTACGAACCTGTAATAAGCGGCCGCGTTTAAACGG

Reverse (PylRS): TTCAGCGGTTTTTTGTCCATGCGGCCGCACCTCCT

Protein sequence (PylRS):
MDKKPLNTLISATGLWMSRTGTIHKIKHHEVSRSKIYIEMACGDHLVVNNSRSSRTARALRHHKYRKTCKRCR
VSDEDLNKFLTKANEDQTSVKVKVVSAPTRTKKAMPKSVARAPKPLENTEAAQAQPSGSKFSPAIPVSTQES
VSVPASVSTSISSISTGATASALVKGNTNPITSMSAPVQASAPALTKSQTDRLEVLLNPKDEISLNSGKPFRELES
ELLSRRKKDLQQIYAEERENYLGKLEREITRFFVDRGFLEIKSPILIPLEYIERMGIDNDTELSKQIFRVDKNFCLRP
MLAPNLANYLRKLDRALPDPIKIFEIGPCYRKESDGKEHLEEFTMLNFCQMGSGCTRENLESIITDFLNHLGIDF
KIVGDSCMVFGDTLDVMHGDLELSSAVVGPIPLDREWGIDKPWIGAGFGLERLLKVKHDFKNIKRAARSESY
YNGISTNL*

#### *WT Ubiquitin plasmid*

WT ubiquitin was used as a control in samples prepared with dispersed TOTAPOL. Here we used the pTXB1-Ub-NpuDnaE-$His_6$ construct described previously.[5] This construct encodes the native ubiquitin sequence, followed by the NpuDnaE intein and a $His_6$ tag. The $His_6$ tag is useful in amber suppression (see below) as it allows the purification of the full-length protein from prematurely truncated polypeptides, while the intein enables the traceless removal of the tag without the use of proteases.[6-7]

*K6X Ubiquitin plasmid*

This construct contained the amber stop codon (TAG) in place of the lysine codon (AAG) at position 6 of the ubiquitin sequence. Site-directed mutagenesis was performed using the NEB Q5 kit, the pTXB1-Ub-NpuDnaE-$His_6$ plasmid and the following primers:

Forward: GATCTTCGTG***TAG***ACTCTGACTG

Reverse: TGCATATGTATATCTCCTTC

Protein sequence:

**MQIFV*TLTGKTITLEVEPSDTIENVKAKIQDKEGIPPDQQRLIFAGKQLEDGRTLSDYNIQKESTLHLVLRLR GG**CLSYETEILTVEYGLLPIGKIVEKRIECTVYSVDNNGNIYTQPVAQWHDRGEQEVFEYCLEDGSLIRATKDH KFMTVDGQMLPIDEIFERELDLMRVDNLPNIKIATRKYLGKQNVYDIGVERDHNFALKNGFIASA*AFNHHHH HH**

*K6X Ubiquitin-$His_6$ plasmid*

This plasmid was prepared from the pTXB1-K6X-Ub-NpuDnaE-$His_6$ construct where the NpuDnaE sequence was deleted using the Q5 mutagenesis kit from New England Biolabs. This produced plasmid pTXB1-K6X-Ub-$His_6$. We used this construct for experiments in bacterial lysates where ubiquitin could be pulled from the lysate if desired using the $His_6$ tag. We removed the intein from the construct to avoid premature intein hydrolysis and to increase the lifetime of the protein in the lysate.

Forward: CATCATCATCATCATCATTAAA

Reverse: ACCACCTCTGAGACG

Sequence:

**MQIFV*TLTGKTITLEVEPSDTIENVKAKIQDKEGIPPDQQRLIFAGKQLEDGRTLSDYNIQKESTLHLVLRLR GG***HHHHHH**

*WT $His_6$-TEV-HP1α plasmid*

The HP1α gene was excised from a GST HP1α plasmid provided by Naoko Tanese[8] (Addgene plasmid # 24074 ; http://n2t.net/addgene:24074 ; RRID:Addgene_24074) and cloned into the pET vector backbone

of a 2BT MacroLab plasmid (generously provided by Dr. Kevin Corbett) using NEBuilder® HiFi DNA Assembly Master Mix. The resulting plasmid contained a His6-TEV-HP1α construct.

Protein sequence:

MKSSHHHHHHENLYFQSGKKTKRTADSSSSEDEEEYVVEKVLDRRVVKGQVEYLLKWKGFSEEHNTWEPEK
NLDCPELISEFMKKYKKMKEGENNKPREKSESNKRKSNFSNSADDIKSKKKREQSNDIARGFERGLEPEKIIGA
TDSCGDLMFLMKWKDTDEADLVLAKEANVKCPQIVIAFYEERLTWHAYPEDAENKEKETAKS*

*$His_6$-TEV-HP1α-2X plasmid*

A TAG codon was inserted at position 2 of HP1α using the NEBuilder® HiFi DNA Assembly Master Mix.

Forward: CAATCCGGA***TAG***AAGAAAACCAAGCGGACA

Reverse: TTTTCTT***CTA***TCCCGGATTGGAAGTACAGGTT

Protein sequence:

MKSSHHHHHHENLYFQSG*KKTKRTADSSSSEDEEEYVVEKVLDRRVVKGQVEYLLKWKGFSEEHNTWEPE
KNLDCPELISEFMKKYKKMKEGENNKPREKSESNKRKSNFSNSADDIKSKKKREQSNDIARGFERGLEPEKIIG
ATDSCGDLMFLMKWKDTDEADLVLAKEANVKCPQIVIAFYEERLTWHAYPEDAENKEKETAKS*

*WT SMC-$His_6$ plasmid*

This construct was purchased from GeneArt Gene Synthesis by ThermoFisher Scientific. The encoding region contains WT Pyrococcus yayanosii SMC (345-468, 694-814) with a SGGS linker and a C-terminal $His_6$ tag. The gene was transferred into the pET vector backbone of the 2BT MacroLab plasmid using NEBuilder® HiFi DNA Assembly Master Mix.

Protein sequence:

MTKGAIVRWGKRKEKLIEEIRAREEERNALVCRLGEIDRTFAVAREEFDTVVKELEEARKSLYEGEARIKRAEEE
KERLKAEILTGEARLPGLRERAENLRRLVEEKRAEISELERRLSSITSKRSGGSLESQSFELRIKLSDLEKELELARKD
LEKVLAEERAVREEIEVAKRRINELDTLIERERGELAKLRGRIERLERKRDKLKKALENPEARELTEKIRAVEKEIAA
LREELSRVEGKLEGLESKGGHHHHHH*

*SMC V6X-His$_6$ plasmid*

Site-directed mutagenesis by HiFi was used to introduce the TAG codon at position 6 of the SMC sequence.

Forward: GCCATT***TAG***CGTTGGGGTAAACGTAAAGA

Reverse: CCCCAACG***CTA***AATGGCACCTTTGGTCATA

Protein sequence:

MTKGAI*RWGKRKEKLIEEIRAREEERNALVCRLGEIDRTFAVAREEFDTVVKELEEARKSLYEGEARIKRAEEE KERLKAEILTGEARLPGLRERAENLRRLVEEKRAEISELERRLSSITSKRSGGSLESQSFELRIKLSDLEKELELARKD LEKVLAEERAVREEIEVAKRRINELDTLIERERGELAKLRGRIERLERKRDKLKKALENPEARELTEKIRAVEKEIAA LREELSRVEGKLEGLESKGGHHHHHH*

**Protein expression and purification**

*K6X Ubiquitin*

The Ub-K6X-NpuDnaE-His$_6$ plasmid was co-transformed with the pULTRA-tRNA$^{Pyl}$/PylRS$^{AF}$ plasmid into BL21 (DE3) competent cells. The cells were grown on an LB agar plate with the presence of ampicillin (100 µg/mL) and spectinomycin (50 µg/mL). After overnight incubation, a colony was selected and inoculated into 5 mL LB media containing antibiotics. After overnight incubation at 37 °C, 1 mL of the preculture was inoculated into 1 L of M9 minimal media supplemented with $^{15}$N-labeled ammonium chloride, $^{13}$C-labeled glucose and antibiotics. The culture was grown at 37 °C with shaking until the $OD_{600}$ reached 0.3 – 0.5. At this point, 10 mL of 0.1 M norbornene lysine stock solution (in 0.5 M NaOH, 1 mM final) was added to the culture and incubated further for 0.5 – 1 h with $OD_{600}$ reaching the 0.5-0.7 range. 1 mL of 0.5 M IPTG (0.5 mM final) was added to the culture and the protein was expressed at 37 °C for 4 hours. After expression, the cells were harvested by centrifugation at 5,000 *×g* and resuspended in 20 mL of cold lysis buffer (50 mM phosphate, 300 mM NaCl, 5 mM imidazole, 1 mM TCEP, Pierce protease inhibitor tablet, pH 7.5). The cells were lysed by sonication and the lysate was clarified by centrifugation at 30,000 *×g*. The supernatant was incubated with 2 mL of Ni-NTA beads for one hour, washed with lysis buffer containing increasing amounts of imidazole (5, 20, and 50 mM imidazole, 20 mL each), and eluted with 12 mL of lysis buffer containing 250 mM imidazole. These steps were performed at 4 °C to avoid premature cleavage of the NpuDnaE intein purification tag. At this point, 120 μL of 1M TCEP (final 10 mM) was added to the elute and incubated at room temperature overnight to cleave the NpuDnaE-$His_6$ intein and to generate ubiquitin without a $His_6$ tag. The sample was then loaded onto a semi-prep C18 column and purified by RP-HPLC. The purified protein was lyophilized and dissolved in 1 mL of 6M guanidine hydrochloride. The solution was dialyzed against 10 mM Tris·KCl buffer (10 mM KCl, pH 7.5) to refold the ubiquitin-norbornene construct. The final sample was analyzed by analytical RP-HPLC and ESI-MS, as well as by solution NMR to confirm the correct fold of the protein (**SI Fig. 2**).

*WT Ubiquitin*

In the case of WT ubiquitin, the pTXB1-Ub-NpuDnaE-$His_6$ plasmid was transformed alone into BL21 (DE3) competent cells. Expression and purification were performed as described above except without the addition of spectinomycin and norbornene lysine.

*HP1α-2X*

BL21(DE3)-Rosetta cells were co-transformed with the HP1α and pUltra plasmids. Pre-cultures were grown at 37 °C overnight under ampicillin and spectinomycin antibiotic selection in LB. Pre-cultures were pelleted in the morning, LB was removed and the pellet was resuspended into $^{13}C$, $^{15}N$ enriched M9 media. Cells were grown at 37 °C until OD600 = 0.8, then induced with 0.5 mM IPTG and 1 mM norbornene lysine. Cells were collected 20 hours later by centrifuging at 5,000 *xg*. The cell pellet was resuspended in lysis buffer (1x PBS, pH 7.4, 300 mM NaCl, 10% glycerol, 7.5 mM imidazole, Roche protease inhibitor tablet) and sonicated at 4 °C. Cell debris were cleared by centrifugation for 30 minutes at 30,000 *xg*. Approximately 5 mL of Ni-NTA resin (per 1 L culture) was added to the lysate and rotated at 4 °C for one hour. The lysate-resin mixture was loaded into a plastic Bio-Rad Econo column and washed with buffer (1x PBS, pH 7.4, 300 mM NaCl, 10% glycerol, 7.5 mM imidazole). Elution buffer (20 mM HEPES, pH 7.2, 300 mM KCl, 400 mM imidazole, 1 mM DTT) was then poured over the column. The eluant was adjusted to 6 M guanidine-hydrochloride and pH 3.0, then applied to a Waters XBridge BEH C18 prep-size RP-HPLC column and fractions were collected using a gradient of 10%-60% solvent B. HP1α eluted around 40% solvent B and due to protein retention on the column, multiple runs were required to collect the protein. The elution product was lyophilized, then refolded by initial resuspension in 6M guanidinium, 20 mM HEPES, pH 7.2, 75 mM KCl and 1 mM TCEP, followed by overnight dialysis against 20 mM HEPES, pH 7.2, 75 mM KCl. Dialysis ensured the removal of TCEP which is necessary prior to reaction with TTz. The final sample was analyzed by analytical RP-HPLC and ESI-MS.

*WT HP1α*

The purification of WT HP1α followed the protocol described above except without the addition of spectinomycin and norbornene-lysine. Additionally, TCEP was kept in the sample to maintain long-term stability.

*SMC-V6X*

Protein expression and cell lysis followed the protocol for HP1α-2X. After clearing the lysate by centrifugation at 30,000 *xg* for 30 min, the lysate supernatant was mixed with Ni-NTA resin, rotated for 1 hr at 4 °C and loaded into a plastic Bio-Rad Econo column. The resin was washed with buffer containing 1x PBS, pH 7.4, 300 mM NaCl, 10% glycerol, 7.5 mM imidazole, and protein elution was performed with buffer containing 20 mM Tris, pH 7.4, 50 mM NaCl, 400 mM imidazole, 1 mM DTT. The elute was adjusted to 6 M guanidine-hydrochloride and pH 3.0, then loaded onto a Waters XBridge BEH C18 prep-size RP-HPLC column. Fractions were collected using a gradient of 10%-60% solvent B (100% ACN + 0.1% TFA). SMC eluted around 40% solvent B. The product was lyophilized, then refolded by initial resuspension in 6M guanidinium, 20 mM Tris, pH 7.4, 50 mM NaCl and 1 mM TCEP. Overnight dialysis against 20 mM Tris, pH 7.4, 50 mM NaCl was performed. Dialysis ensured the removal of TCEP which is necessary prior to reaction with TTz. The final sample was analyzed by analytical RP-HPLC and ESI-MS.

*WT SMC*

The purification of WT SMC followed the protocol described above except without the addition of spectinomycin and norbornene-lysine.

### Conjugation of TTz to proteins

*Ub-TTz*

Lyophilized K6X ubiquitin was dissolved in 1 mL of 6 M guanidine hydrochloride buffer (10 mM Tris-HCl, 10 mM KCl) to a final concentration of 100 μM. TTz was dissolved in DMSO to prepare a 5 mM stock solution and 100 μL of the stock were added to the ubiquitin solution for a final TTz concentration of 500 μM. The mixture was mixed gently overnight and the completion of the reaction was confirmed by the absence of a K6X ubiquitin peak in the analytical RP-HPLC trace (**SI Fig. 3**). The solution was then dialyzed against 10 mM Tris, 10 mM KCl, pH 7.5 buffer to refold the protein and remove excess TTz. The complete removal of unbound TTz was confirmed by analytical RP-HPLC and the final conjugated Ub-TTz product was confirmed by ESI-MS (**SI Fig. 4**). Since tetrazine-norbornene conjugates have absorbance at A280, we determined protein concentration in the following way. Serial dilutions of WT ubiquitin and Ub-TTz were loaded on an SDS-PAGE gel and the density of the bands and the A280 absorbance of the samples were compared. Thus, it was determined that the same amount of Ub-TTz has 8-fold higher absorbance at A280 compared to WT ubiquitin, producing an extinction coefficient of 10,240 $cm^{-1}\cdot M^{-1}$ for Ub-TTz vs 1280 $cm^{-1}\cdot M^{-1}$ for WT ubiquitin.

*HP1α-TTz*

Refolded HP1α-2X was dialyzed against 20 mM HEPES, pH 7.2, 75 mM KCl and diluted to 20 μM. 200 μM TTz from a 50 mM stock was added to the HP1α solution. The mixture was gently rotated for one hour until reaction completion was confirmed by the retention time shift of the HP1α-2X RP-HPLC peak (**SI Fig. 3)**. The solution was then concentrated to both concentrate HP1α and remove excess TTz. We found that TTz is completely removed after several buffer exchanges (**SI Fig. 3 & 4**). The final concentration of HP1α-TTz was determined by SDS-PAGE gel analysis as performed for Ub-TTz.

*SMC-TTz*

Refolded SMC-V6X was dialyzed against 20 mM Tris, pH 7.4, 50 mM NaCl and diluted to 25 μM. 250 μM TTz from a 50 mM stock was added to the SMC solution. The mixture was gently rotated for three hours until reaction completion was confirmed by the retention time shift of the SMC-V6X RP-HPLC peak (**SI Fig. 3)**. The solution was then concentrated to both concentrate SMC and remove excess TTz (**SI Fig. 3 & 4**). The final concentration of SMC-TTz was determined by SDS-PAGE gel analysis as done performed Ub-TTz.

**DNP sample preparation**

*Ub-TTz*

Refolded Ub-TTz in 10 mM Tris, 10 mM KCl was concentrated and the protonated buffer was exchanged several times with deuterated buffer with the same composition (10 mM Tris-$d_{11}$, 10 mM KCl, pD 7.5 in $D_2O$). The final concentration of protein was 15 mg/mL (1.66 mM) and 97% deuteration of the solvent was achieved ($D_2O$:$H_2O$ = 97:3). The sample was diluted with deuterated buffer to the desired protein concentration and 9 µL of the sample were combined with 1 µL of glycerol-$d_8$, and transferred into a 1.9 mm DNP zirconia rotor.

*WT ubiquitin with 15 mM TOTAPOL*

Refolded WT ubiquitin in 10 mM Tris, 10 mM KCl, pH 7.5 was concentrated and the protonated buffer was exchanged several times with 60% glycerol-$d_8$ in 90% deuterated buffer with the same composition (glycerol-$d_8$:$D_2O$:$H_2O$ = 60:4:36). The **f**inal concentration of protein was 11.2 mg/mL (1.28 mM). The sample was diluted with the deuterated buffer (final 10.5 mg/mL) and 9.5 µL of the sample were combined with 0.5 µL of 300 mM TOTAPOL in 90% deuterated buffer with the same composition, and transferred into a 1.9 mm DNP zirconia rotor.

*HP1α-TTz*

HP1α-TTz was concentrated and the protonated buffer was exchanged multiple times into 20 mM HEPES, 50 mM KCl, pD 7.2 in 97% $D_2O$. The final concentration of protein was 200 μM. The sample was then diluted to 50 μM in deuterated buffer and 10% ($^{13}$C-depleted 0.05%) glycerol-$d_8$ and transferred into a 1.9 mm DNP zirconia rotor.

*SMC-TTz*

SMC-TTz was concentrated and the protonated buffer was exchanged multiple times into 20 mM Tris, 50 mM NaCl, pD 7.4 in 97% $D_2O$. The final concentration of protein was 250 μM. The sample was diluted to 50 μM in deuterated buffer and 10% ($^{13}$C-depleted 0.05%) glycerol-$d_8$ and transferred into a 1.9 mm DNP zirconia rotor.

**Solution NMR experiments**

Experiments were performed using a Bruker Avance III 600 MHz ($^1$H Larmor frequency) NMR spectrometer equipped with a triple resonance probe. Chemical shifts were referenced to the TMS frequency. The HSQC spectra were acquired using the standard Bruker fhsqcf3gpph pulse sequence. The HSQC spectrum was acquired with 16 scans while using the following parameters: 298 K, 2048 points and 122 ms acquisition in the direct dimension, 256 points and 58 ms acquisition in the indirect dimension, interscan delay of 1.5 s, $^1$H center frequency of 4.7 ppm, $^{15}$N center frequency of 118 ppm.

**MAS DNP NMR experiments**

Experiments were performed on a 600 MHz Bruker DNP NMR spectrometer equipped with a NEO console and a 395 GHz gyrotron for high-power microwave irradiation. All spectra were recorded using a triple resonance ($^1$H, $^{13}$C, $^{15}$N) 1.9 mm MAS Bruker probe. The magic angle was set with KBr and the $^{13}$C chemical shift was referenced to adamantane[9] (40.49 ppm) at room temperature or the rotor silicon plug $^{13}$C peak (3.2 ppm) at low temperature. Sample temperature was maintained at 100 K, at either 12 or 24 kHz. Temperature was confirmed by the T1 of a KBr standard.[10] The largest signal enhancements were recorded at microwave power of 5 W.

1D $^1$H-$^{13}$C CP experiments for enhancement determination had the following parameters: 100 K, 12 or 24 kHz MAS, 5 s interscan delay, 20 ms acquisition, 2048 points, $^{13}$C center frequency set at 100 ppm. Scans were tailored for each sample to achieve sufficient signal-to-noise in the microwave off spectrum. For CP, a 500 μs contact time was used with the Hartmann-Hahn matching condition satisfied by setting the $^{13}$C power to 66 kHz and optimizing on the $^{1}$H channel. 100 kHz spinal64 decoupling[11] was performed during acquisition. DNP build-ups were measured with this 1D $^{1}$H-$^{13}$C CP experiment but with varied interscan delays until saturation.

The 2D $^{13}$C -$^{13}$C CORD experiment used the CORDxy4 pulse sequence[12], with the following parameters: 100 K, 24 kHz MAS, 64 scans, 5 s interscan delay, 1536 points and 15 ms acquisition in the direct dimension, 288 points and 4 ms acquisition in the indirect dimension, $^{13}$C center frequency of 100 ppm. The CORD conditions were set to 24 kHz ($1*\omega_r$) and 12 kHz ($0.5*\omega_r$). For CP, a 500 μs contact time was used, together with 50 ms of CORD mixing. 100 kHz spinal64 decoupling was performed during acquisition. To process the spectrum, a GM window function was used with 75 Hz line broadening in each dimension.

**DNP of bacterial lysates**

*Sample preparation*

The K6X Ubiquitin-His$_6$ plasmid and the pULTRA-tRNA$^{Pyl}$/PylRS$^{AF}$ plasmid were co-transformed into BL21 (DE3) competent cells. We followed the expression protocol for K6X ubiquitin as described above with the exception that we only used 200 mL of M9 media enriched with $^{13}$C-glucose and $^{15}$N-labeled ammonium chloride. After the end of the expression period, the cells were lysed by sonication in 5 mL lysis buffer containing 50 mM phosphate, 300 mM NaCl, 5 mM imidazole, 1 mM TCEP, Pierce protease inhibitor tablet, pH 7.5. After lysis, the solution was dialyzed overnight against buffer containing 20 mM phosphate and 150 mM NaCl, pH 7.2. This step was performed to remove endogenous thiol molecules as they can reduce the lifetime of nitroxide-based polarization agents.[13] At this point, 50 uL of 100 mM TTz in DMSO (final 1 mM) were added to the dialyzed solution and the mixture was mixed gently. After 8 hours, 50 μL of the reaction mixture was removed and diluted 20 times in deuterated phosphate buffer with the same salt composition as above. The diluted solution was concentrated using a 3,000 MWCO concentration device and the buffer was exchanged several times to achieve 95% deuteration level. Complete removal of unbound TTz was confirmed by the disappearance of the TTz peak in the analytical RP-HPLC trace. To prepare the sample for DNP, 9 μL of the sample were combined with 1 μL of glycerol-$d_8$ and the solution was transferred into a 1.9 mm zirconia rotor.

To determine the amount of ubiquitin in the lysate, we resolved the protein content of the lysate sample by SDS-PAGE gel electrophoresis (**SI Fig. 9**). The intensities of all visible bands were quantified using the ImageJ analysis software and the intensity of the ubiquitin band was divided by the combined intensity of all the other protein bands in the gel, resulting in ~5% ubiquitin content. Furthermore, the intensity of the ubiquitin band was compared to the intensities of known amounts of purified ubiquitin loaded on the same gel. This gave an estimate of the absolute amount of ubiquitin in the lysate DNP sample, which was ~ 50 μg.

Finally, to determine the efficiency of the TTz-norbornene reaction in this context, we used Ni-NTA affinity column purification to remove ubiquitin from the lysate. The purified ubiquitin was then subjected to LC-MS, confirming the presence of substantial amounts of Ub-TTz (**SI Fig. 9**).

*Analysis of the DNP signal enhancements*

One-dimensional $^1H$-$^{13}C$ cross-polarization experiments were acquired with and without microwaves. After applying line broadening of 100 Hz and polynomial baseline correction, the signal intensity at the largest Cα peak (54.4 ppm for Ub, 57.4 ppm for SMC, 54.7 ppm for HP1α) was measured for the on and off states. The on to off ratio is reported as the enhancement ε .

*Quantification of the selective DNP enhancements in lysates*

To calculate the enhancement of Ub-TTz in lysates, we followed a method previously described by Viennet et al.[14] We began by measuring the peak intensity at 54.4 ppm for the ON signal (6.11 x $10^9$) and OFF signal (2.44 x $10^9$). The OFF signal was then multiplied by two (4.88 x $10^9$) to match the intensity of the glycerol signals from each spectrum. The glycerol peaks arise from the sample solvent and we assume that their enhancement represents the non-selective enhancement of the lysate background. The difference between the ON signal and 2x the OFF signal was 1.22 x $10^9$ and represents the enhancement on ubiquitin. Since ubiquitin is only 5% of the total protein in the sample, we assumed that only 5% of the microwave OFF signal is due to ubiquitin (1.22 x $10^8$). Therefore, the "true" enhancement of ubiquitin in the sample is the ratio of 1.22 x $10^9$ and

$1.22 \times 10^8$ , i.e. it is equal to 10. We divided this by two to account for the two-fold enhancement on the background glycerol signals, which then produces a selective ubiquitin enhancement equal to 5.

**Electron paramagnetic resonance spectroscopy**

EPR spectra were acquired on a 9 GHz EMX Bruker EPR spectrometer at room temperature. We recorded the EPR spectra on samples taken directly from the DNP NMR rotors and placed in quartz capillaries. The sample length in the capillary spanned the whole length of the microwave cavity so that the concentrations of radicals in all samples could be compared to a 50 and 100 µM TOTAPOL standards.

**Calculation of the distance between polarization agents at various concentrations**

We followed the application of the Wigner-Seitz radius as done previously by Pinon et al.[15]

$$d = 2\left(\frac{3}{4\pi C N_A}\right)^{1/3}$$

Here $d$ is distance between the biradical polarization agents, $C$ is the concentration of polarization agents, $N_A$ is Avogadro's number.

**Analysis of the polarization buildup curves**

Peak intensities at the 54.4 ppm Cα peak were measured using TopSpin 4.0.5. Using GraphPad Prism version 8.2, data were fit to either a monoexponential association model, $I = I_0 + \left(1 - e^{-\frac{t}{\tau}}\right)$ or a biexponential association model, $I = I_0 + a\left(1 - e^{-\frac{t}{\tau_1}}\right) + b\left(1 - e^{-\frac{t}{\tau_2}}\right)$ where $a$ and $b$ are fit parameters that account for the fraction of fast and slow components and $\tau_1$ and $\tau_2$ are the build-up time constants of those components. Here $I_0$ = 0.

### Supplementary figures

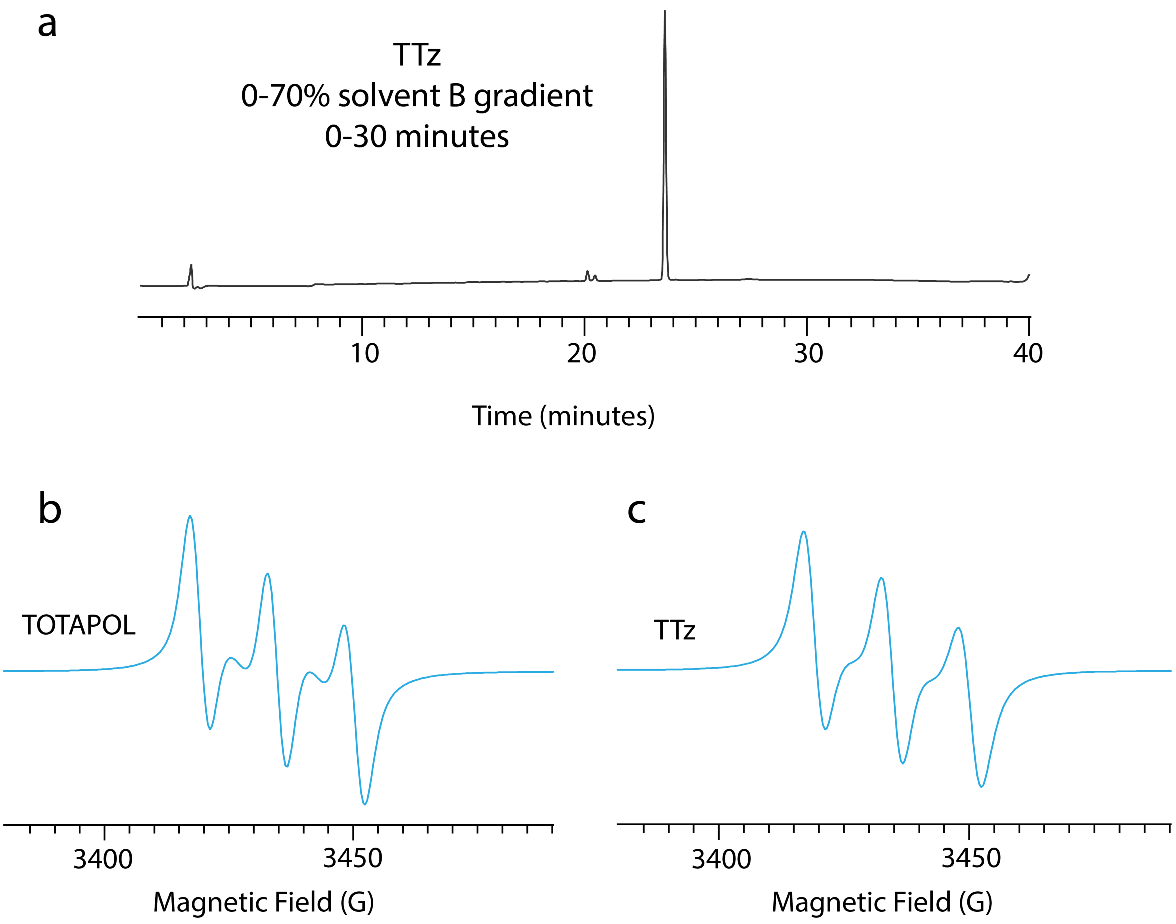


**SI Figure 1:** Analysis of the TOTAPOL-tetrazine (TTz) polarization agent. (a) RP-HPLC trace at 214 nm absorbance of purified TTz. (b) A 9 GHz solution EPR spectrum of TOTAPOL, and c) TTz. Spectra were acquired in DMSO-$d_6$. TTz displays the characteristic nitroxide spectral features.

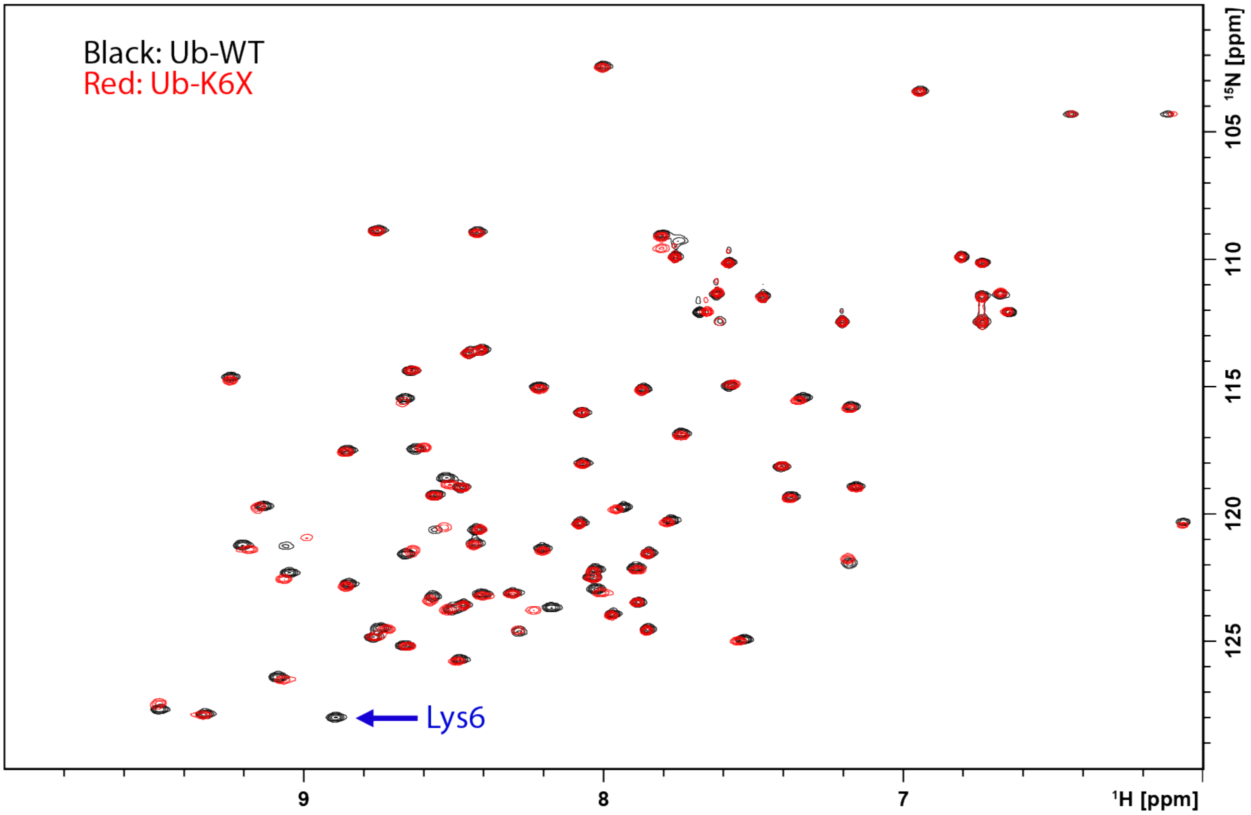


**SI Figure 2:** $^{1}$H-$^{15}$N HSQC spectra of wild-type ubiquitin (black) and ubiquitin incorporating norbornene-lysine at position 6 (Ub-K6X) (red). Since the UAA is supplied at natural abundance, the Lys6 cross-peak is missing from the spectrum. The UAA causes only minor local perturbations of the ubiquitin fold.

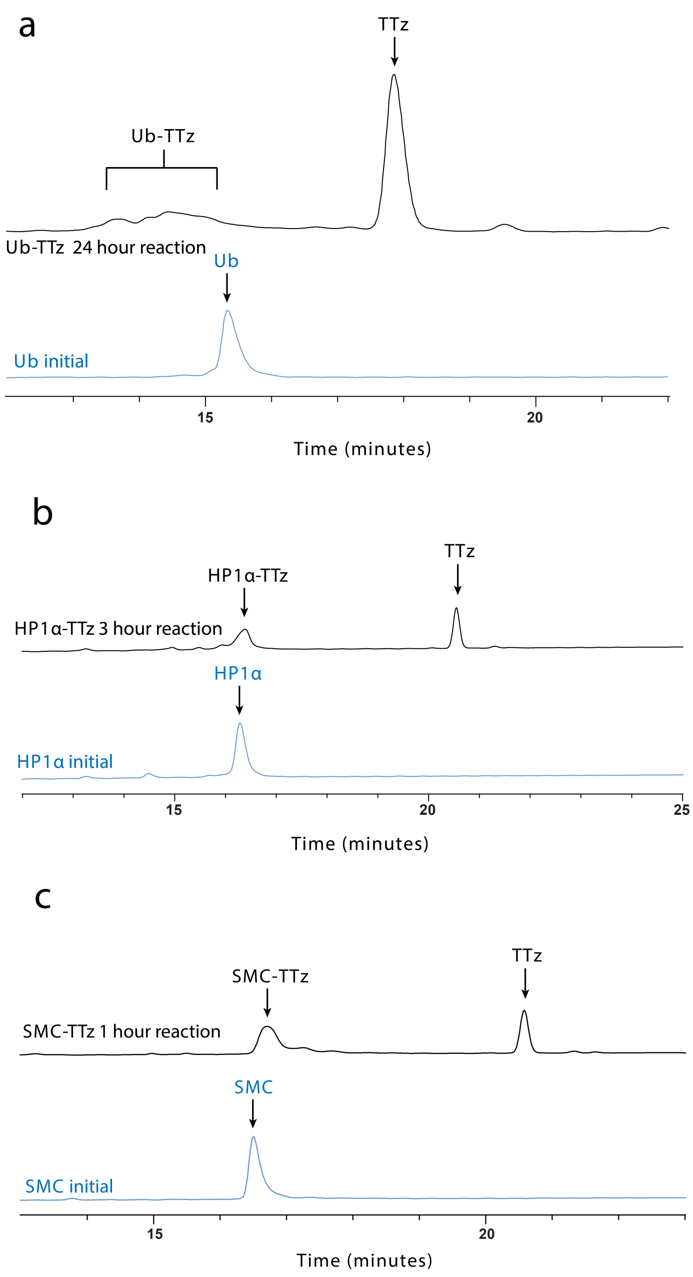


**SI Figure 3:** Protein-TTz reactions monitored by analytical RP-HPLC for (a) Ub-TTz, (b) HP1α -TTz, and (c) SMC-TTz. The conjugation reaction results in a shift of the retention time and peak broadening due to multiple isomer products with the same mass. Unreacted TTz has strong absorbance at 214 nm and 280 nm, which was useful in monitoring TTz removal from the final DNP samples.

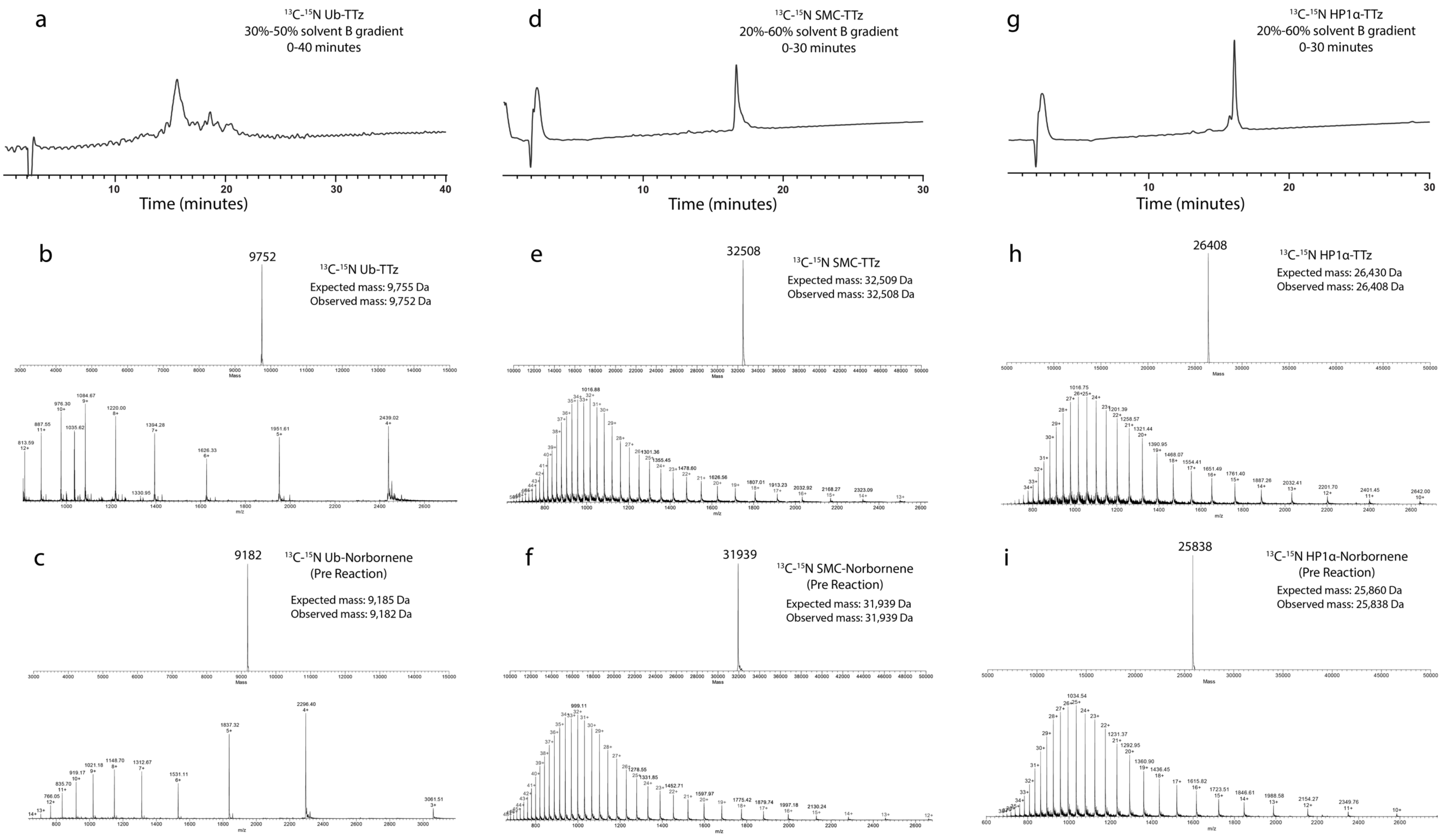


**SI Figure 4:** Purified $^{13}$C,$^{15}$N-labeled protein-TTz samples analyzed by RP-HPLC and ESI-TOF-MS. (a), (d) and (g) show the RP-HPLC traces for purified Ub-TTz, SMC-TTz and HP1α-TTz respectively. (b), (e), and (h) show the ESI-MS of the pure conjugated proteins, while (c), (f) and (i) show the ESI-MS of the proteins before TTz conjugation.

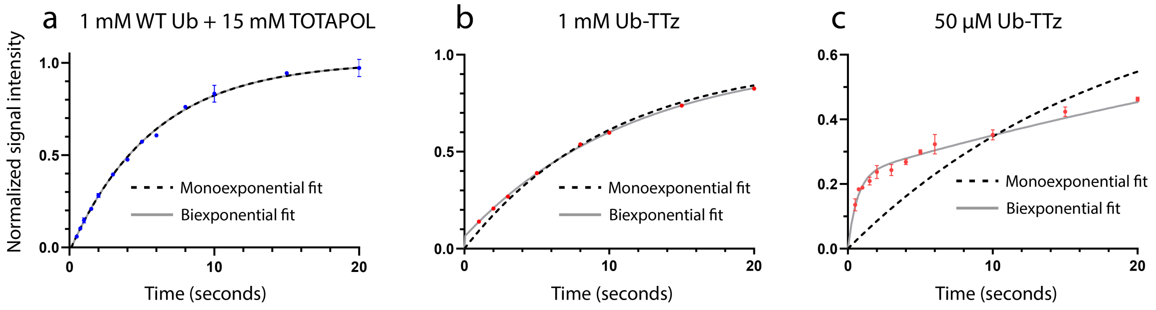


**SI Figure 5:** DNP build-up curves identical to Figure 4 in the main text with zoomed axes for (a) 1 mM WT Ub + 15 mM TOTAPOL , (b) 1 mM Ub-TTz, and (c) 50 µM Ub-TTz. At this scale, the initial build-up clearly depicts a bimodal increase in the DNP signal for 50 µM Ub-TTz. Error bars correspond to the standard deviation of two independent measurements of the enhancements.

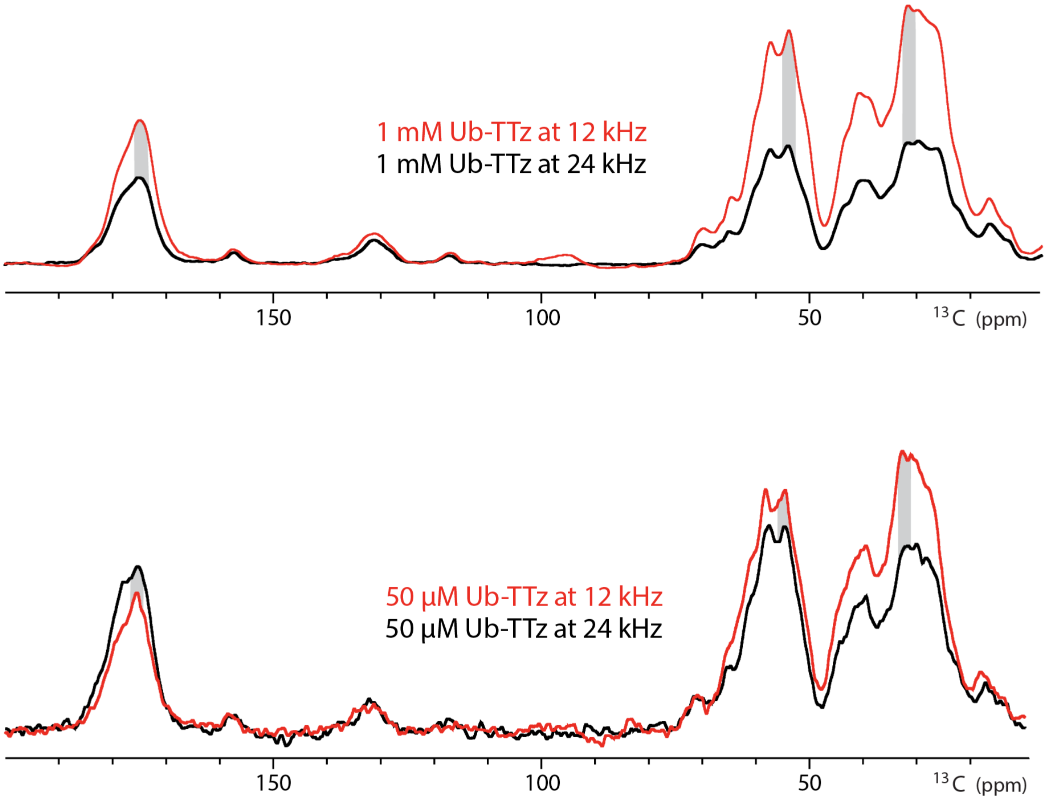


MAS dependence based on MW ON signal intensity

| Sample | Ratio of signal intensity 24 kHz : 12 kHz | | |
|---|---|---|---|
| | Cβ, Cγ (31 ppm) | Cα (54 ppm) | C′ (175 ppm) |
| 50 µM Ub-TTz | 0.69 | 0.85 | 1.23 |
| 1 mM Ub-TTz | 0.47 | 0.51 | 0.61 |
| 1 mM WT Ub + 15 mM TOTAPOL | 0.46 | 0.56 | 0.78 |

**SI Figure 6:** Comparison of the DNP-enhanced (microwave on) signal intensities as a function of MAS frequency for 1 mM Ub-TTz (top) and 50 µM Ub-TTz (bottom). The table summarizes the ratio of signal intensity at 24 kHz to 12 kHz MAS for different regions of the spectrum.

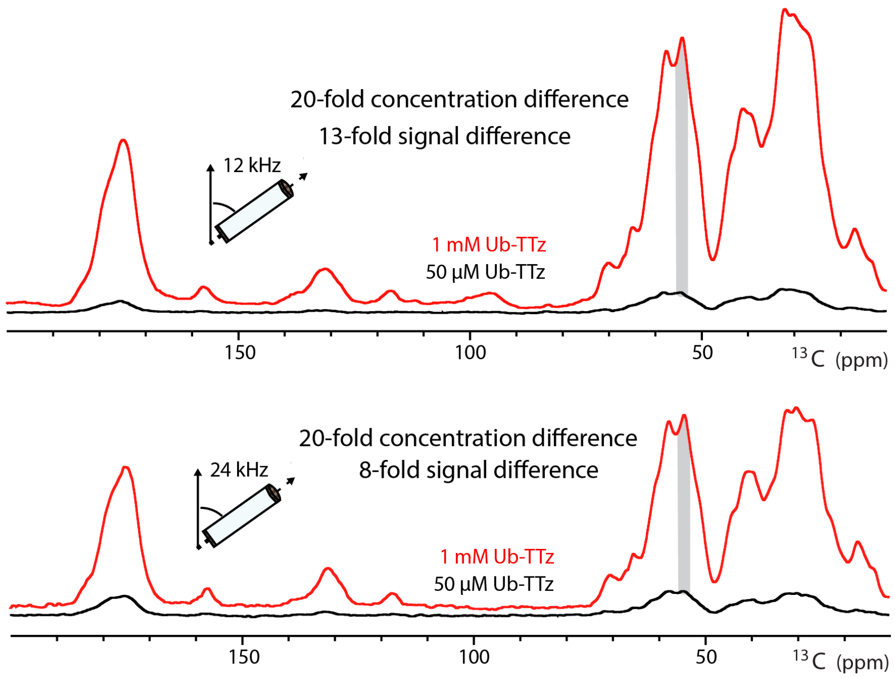


Comparative MAS dependence based on MW ON signal intensity

| Samples | Fold difference Ub concentration | Fold difference MW ON @ 12 kHz MAS | Fold difference MW ON @ 24 kHz MAS |
|---|---|---|---|
| 1 mM Ub-TTz : 50 μM Ub-TTz | 20 | 13.15 | 7.90 |
| 1 mM WT Ub : 50 μM Ub-TTz | 20 | 13.81 | 9.02 |
| 1 mM WT Ub : 1 mM Ub-TTz | – | 1.05 | 1.1 |

**SI Figure 7:** Comparison of absolute enhanced signal intensities for samples containing different concentrations of ubiquitin-TTz or ubiquitin with dispersed TOTAPOL. The purpose of this comparison is to illustrate that smaller microwave on/off enhancements do not necessarily translate to less NMR signal. For example, the 20-fold concentration difference between the 50 μM and 1 mM Ub-TTz samples would imply that a 20-fold difference in NMR signal intensities is to be expected. However, the measured absolute signal difference is much smaller and decreases at higher MAS. The comparison also shows nearly identical signal intensities for the 1 mM Ub-TTz sample and the 1 mM ubiquitin with 15 mM dispersed TOTAPOL sample, despite their very different enhancements. We attribute this effect to radical-induced depolarization in the absence of microwaves which is expected to be more prominent in samples containing higher radical concentrations. Depolarization leads to artificially lower signals in the microwave off spectra and thus skews the reported on/off enhancements.

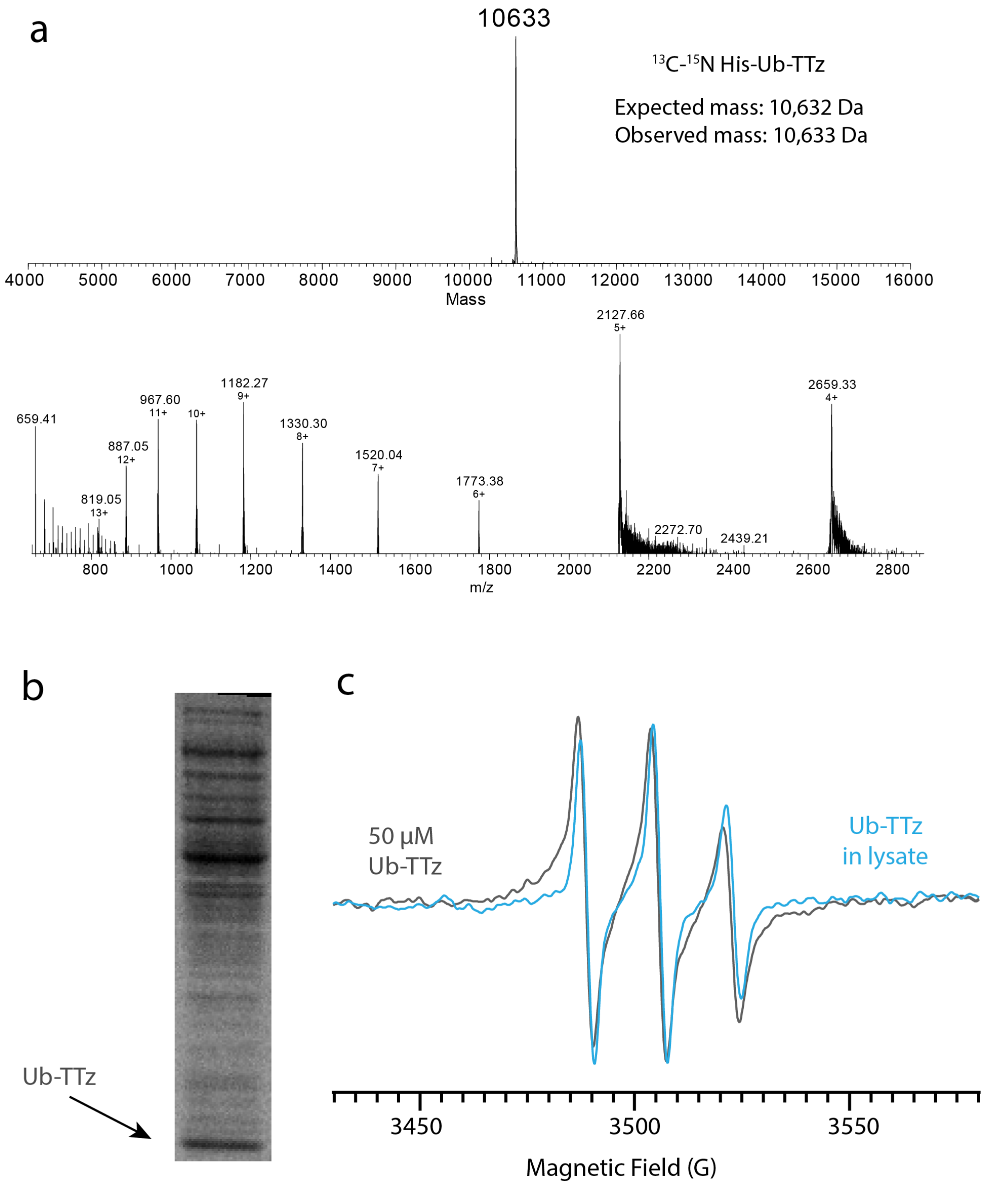


**SI Figure 8:** Analysis of Ub-TTz in lysate samples. Ub-TTz in lysates contained a $His_6$ tag enabling the affinity column purification of the protein from the lysate and the LC-MS analysis of the conjugation efficiency. (a) ESI-TOF MS of Ub-TTz purified from the lysate. (b) SDS-PAGE analysis of the lysate sample. Analysis of the protein band intensities allowed us to quantify the percentage of ubiquitin present in the lysate (5% of the total protein content and present at 40 µM). (c) 9 GHz EPR of the lysate sample confirmed that Ub-TTz is present at the expected 40 µM concentration.

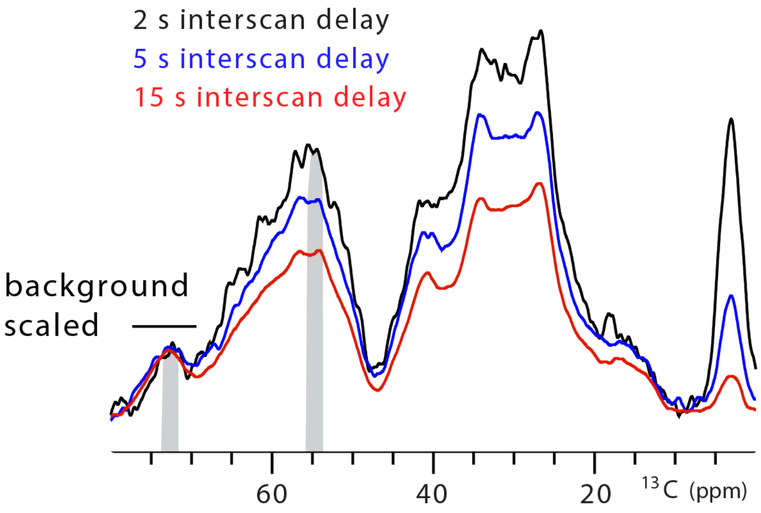


Ub-TTz selectivity variation by interscan delay

| Sample | Ratio of signal intensity Cα : glycerol | | |
|---|---|---|---|
| | 2 s | 5 s | 15 s |
| Ub-TTz in lysate | 4.03 | 3.47 | 2.59 |

**SI Figure 9:** The selective DNP enhancement of ubiquitin in lysates can be tuned as a function of the interscan delay. Overlaid spectra were acquired with 256 scans and interscan delays of 2 seconds, 5 seconds and 15 seconds. The table summarizes the ratio of protein signal (Cα) to the background glycerol signals.

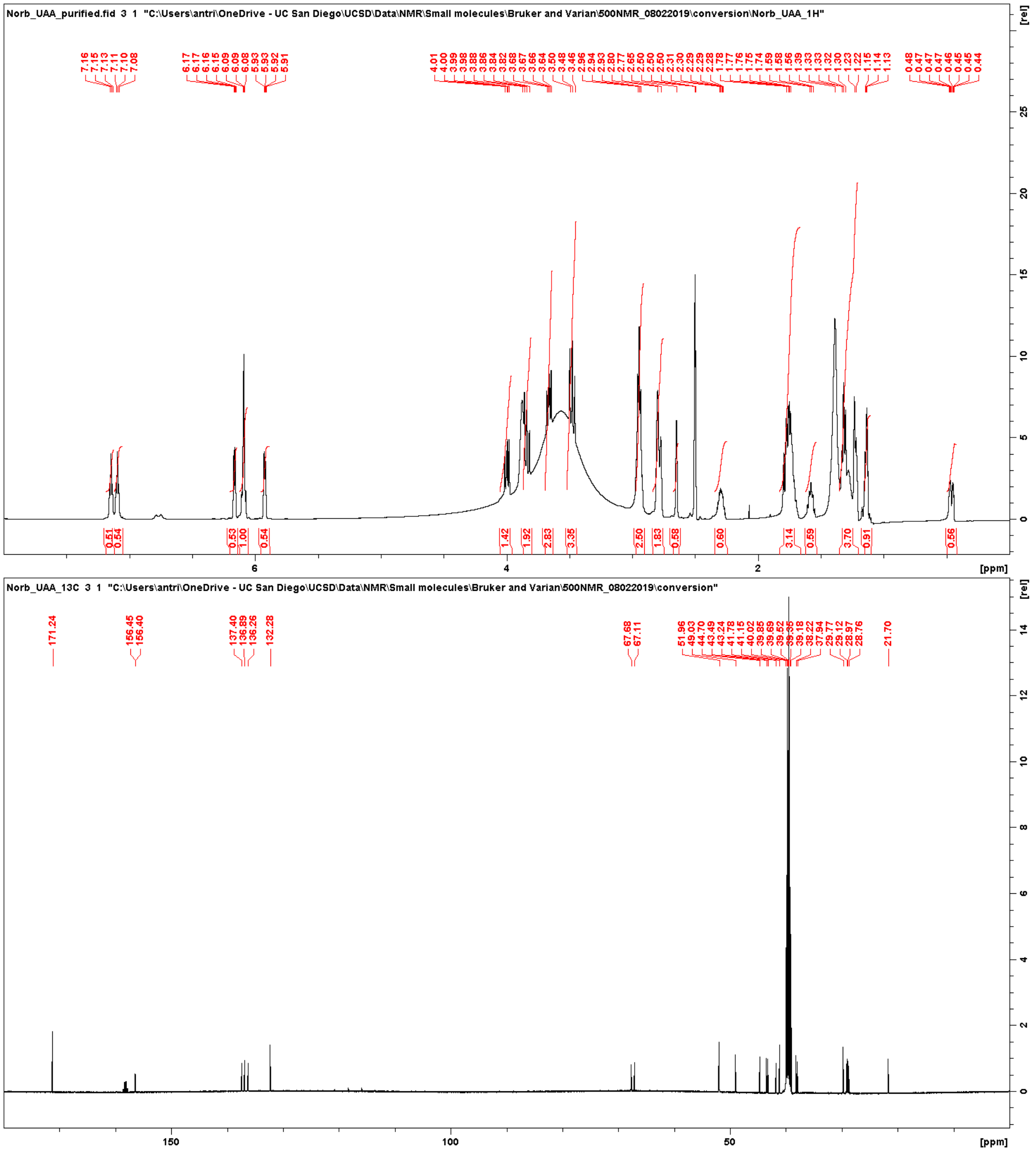


**SI Figure 10:** $^{1}$H and $^{13}$C NMR spectra of norbornene-lysine.

Enhancements

| Sample | Concentration | 12 kHz MAS (ε) | | 24 kHz MAS (ε) |
|---|---|---|---|---|
| Ub-TTz | 50 µM | 11 (1.5 s ID) | 9 (5 s ID) | 11 |
| | 1 mM | 24 | | 18 |
| WT Ub + 15 mM TOTAPOL | 1 mM | 35 | | 22 |
| SMC-TTz | 65 µM | 7 | | – |
| HP1α-TTz | 50 µM | 6 | | – |

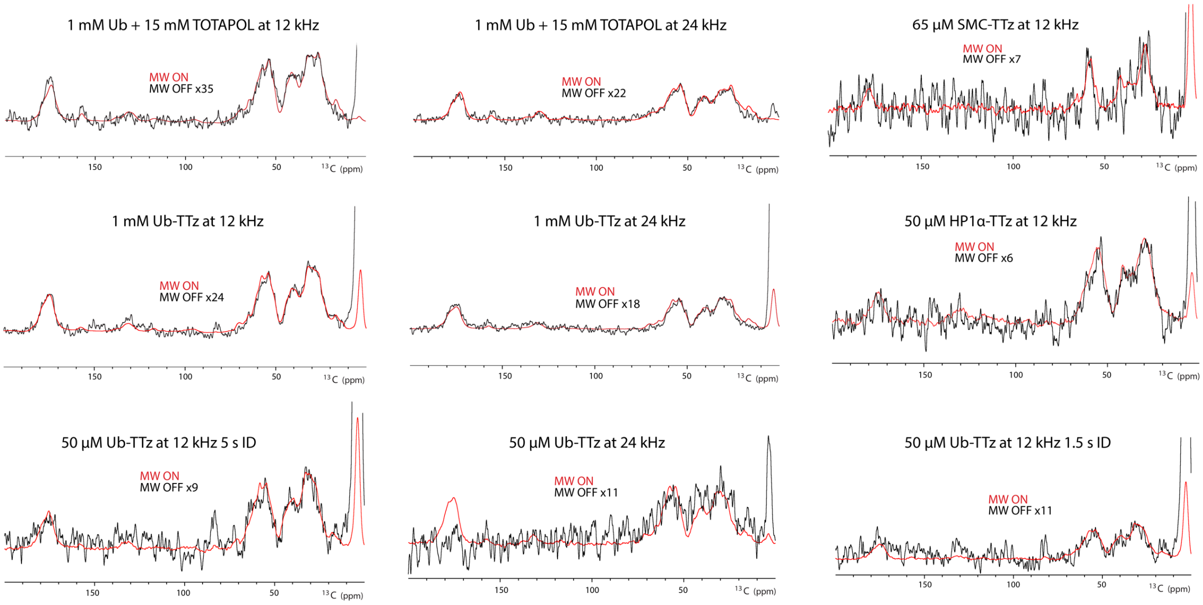


**SI Figure 11:** Summary of the raw data and experimental parameters used to measure the DNP enhancements reported throughout the manuscript. ID = interscan (recycle) delay.
